# Bridging the gap between omics and structural data: A framework for interpreting protein-RNA interaction specificity

**DOI:** 10.64898/2026.09.24.754079

**Authors:** Yoann Fauconnet, Shuyun Deng, Rebecca Koundri, Martin Pagani, Chloé Quignot, Déborah Antonio, Jessica Andreani

## Abstract

Protein-RNA interactions play a central role in many cellular processes, such as gene regulation, protein synthesis and viral infections. Although a few thousand protein-RNA complexes have been structurally characterized, they represent only a small fraction of all interactions. This lack of experimental data limits the ability of deep-learning based approaches such as AlphaFold3 to predict structures of protein-RNA interactions. In this study, we investigate protein-RNA binding specificity by enriching experimental structures with omics data. To this end, we present a scoring approach and associated web resource quantifying the agreement between experimental structures of protein-RNA interfaces and their omics-derived binding preferences, allowing us to identify interaction motif cores. Through key structural and evolutionary features, we further highlight that these motif cores correspond to important interface regions. We then leverage the dataset of protein-RNA complexes for which structural information can be combined with binding preferences in order to benchmark the ability of AlphaFold3 to predict protein-RNA interaction specificity. To this aim, we run AlphaFold3 predictions using different RNA inputs, from the exact sequence present in the experimental structure to a non-specific sequence, including sequences embedding the consensus binding motif from *in vitro* experiments. We show that despite good prediction quality, the sensitivity of AlphaFold3 to the exact RNA sequence used as input indicates signs of memorization. We examine some particular complexes and uncover challenges encountered by our scoring workflow and AlphaFold3, especially in the case of alternative binding modes. Altogether, this work evidences both the promise and current limitations of deep learning approaches for protein-RNA structure prediction, and provides a resource to guide their further development.

**AUTHOR SUMMARY:** Proteins and RNA interact constantly within cells, driving processes such as gene expression and protein synthesis. Understanding the mechanisms underlying specific protein-RNA recognition matters for both biological knowledge and therapeutic applications, yet remains difficult. Far fewer experimentally resolved structures exist for protein-RNA complexes than for proteins alone, and structure prediction tools based on artificial intelligence, such as AlphaFold3, still struggle with RNA and protein-RNA interactions. In this study, we combine experimental data from different sources to design a scoring method quantifying their agreement and we show our method’s ability to highlight relevant binding regions. These results are available for interactive exploration through a web resource. We also benchmark AlphaFold3 for its ability to predict protein-RNA interactions in a specific manner. This benchmark highlights difficulties of AlphaFold3 in producing specific predictions and signs of memorization. Overall, this work showcases the interest of combining different data modalities and points to possible guidelines for future improvement of specific structural predictions of protein-RNA interactions.

## INTRODUCTION

Protein-RNA interactions (PRIs) are central to a myriad of cellular processes, orchestrating gene expression, post-transcriptional regulation, and protein synthesis [1]. These interactions are fundamental to cellular homeostasis, and their dysregulation is implicated in a wide spectrum of human diseases, such as cancers, neurodegenerative disorders, viral infections, and autoimmune diseases [2–4]. A comprehensive understanding of these interactions is therefore critical for deciphering complex biological pathways and developing targeted therapeutic solutions. Despite their pivotal roles, the intricate mechanisms governing PRIs remain largely enigmatic. Elucidating these mechanisms necessitates detailed structural insights, traditionally obtained through experimental techniques such as X-ray crystallography, nuclear magnetic resonance (NMR) spectroscopy, and cryo-electron microscopy (cryo-EM) [5]. However, out of more than 250,000 entries in the Protein Data Bank (PDB) [6, 7] as of 2026, only a few thousand actually contain interacting protein and RNA molecules. This collection represents a limited and largely redundant subset of the vast landscape of PRIs.

Early computational methods to predict protein-RNA interface structure were mostly template-based [8, 9], although several template-free and hybrid docking methods developed for protein-protein interactions were extended to protein-RNA interactions [10, 11], albeit with success limited by the conformational flexibility of both protein and RNA in these interfaces [12]. Consistently with the functional importance of PRIs, translating into selection pressure, several studies, including a recent study from our team, have demonstrated that interactions between homologous proteins and RNAs across different species are often structurally conserved, even between remote homologs [13, 14]. Additionally, evolutionary signals in both proteins [15] and RNA [16] can provide important information about their interactions.

The rise of deep learning-based approaches powered by evolutionary information, such as AlphaFold, has ushered in a new era of structural biology [17]. The most recent of these methods, in particular AlphaFold3 (AF3) [18], offer the potential to significantly expand the repertoire of protein-RNA complex structures. However, they currently exhibit limitations in predictive accuracy for protein-RNA interactions, with reported success rates hovering around 40% for AF3, much lower than the 70% success rate for protein-protein interaction structural prediction [18, 19]. The limited availability of high-quality experimental structures for training hinders the development of robust predictive approaches. Furthermore, PRIs present a complex challenge for structural prediction compared to protein-protein interactions due to several properties, such as their flexibility and their diverse specificities, ranging from strict sequence dependence to structural recognition to broadly non-specific interactions [20]. This underscores the ongoing need for refined computational approaches and complementary experimental investigations to fully unravel the complexity of PRIs.

As a means to mitigate the lack of structural data about PRIs, we set out to enrich protein-RNA data by integrating information from other experimental methods. We built a dataset incorporating RNA-binding preferences obtained through high-throughput in vitro and in vivo experiments. We introduced an original agreement scoring workflow to assess consistency between these data sources, enabling filtering of interactions and identification of high-confidence PRIs. By then integrating evolutionary information with structural and high-throughput data, we gained a more comprehensive view of the determinants of PRI specificity.

We also used the dataset of protein–RNA complexes with both experimental structures and experimental binding preferences available to assess the ability of AF3 to predict and score specific protein-RNA interfaces. Although the dataset is largely redundant with the AF3 training data, comparing structural predictions involving different RNA sequences enabled us to highlight the need to exercise caution when interpreting the prediction specificity and confidence metrics. To further illustrate the challenges of data integration and structural prediction of PRIs, we investigated two selected PRIs from our dataset in deeper detail, using our agreement score as a key analytical tool.

## RESULTS

### 1. Agreement scoring captures RNA-binding specificity from structural and high-throughput RNA-binding assay data

The limited number of experimentally determined protein-RNA interface structures represents a limitation both for understanding these interactions in molecular detail and for training reliable structure prediction methods. In this study, we sought to explore how detailed structural data can be enriched by binding specificity information derived from high-throughput RNA-binding assays. How to integrate the two data modalities represented a challenge, because the two approaches provide complementary but very different views of RNA recognition. Structural data describe molecular contacts within a specific protein–RNA complex, whereas high-throughput RNA-binding assays capture sequence preferences, typically represented as ranked lists of short sequences. In this study, we developed a scoring workflow to assess the agreement between RNA-binding profiles derived from experimental sequence preferences and RNA sequences observed in interface 3D structures. We found that a two-step alignment procedure, in which top-ranked sequences first define a robust profile and additional sequences subsequently enrich it, provided a robust representation of the binding preferences (see Methods). Position-specific mono- and dinucleotide frequency matrices were then derived from these alignments and used for agreement scoring.

We applied this framework to a dataset of human protein–RNA interactions (PRIs) for which both experimental structures and independent RNA-binding data were available. The latter comprised RNAcompete (RNAC) [21] and RNA Bind-n-Seq (RBNS) [22], which measure protein–RNA binding preferences *in vitro*, and Cross-linking followed by immunoprecipitation (CLIP) [23], which captures protein–RNA interactions *in vivo* (see Methods). CLIP was used as complementary evidence to the *in vitro* assays, and proteins featuring only CLIP data were excluded. For this purpose, we built a dataset of human protein-RNA interactions (PRI) with both available experimental PRI structural data from the PDB and RNA binding preference data from RNAC, RBNS and CLIP experiments (see Methods). After filtering to remove short proteins and small interfaces (see Methods), we obtained a dataset of 294 PRIs involving 60 proteins with RNAC, RBNS and CLIP data available for 38, 37 and 14 proteins, respectively (Fig S1).

We developed a workflow to compute an agreement score S reflecting the concordance between structural data and RNA binding preferences deduced from omics data (Fig 1). By design, the values of S are negative, higher values reflect better agreement, and S=0 indicates perfect agreement. Concretely, this score measures how closely the RNA sequence bound to the protein in the 3D structure matches the experimentally identified RNA 5-mers (from RNAC, RBNS or CLIP data) that interact with that same protein (see Methods). This scoring is dinucleotide-based to capture precise interaction specificity as highlighted in previous studies [24] and was also employed to identify the sub-region (dubbed “motif core”) within the structural protein-binding RNA sequence exhibiting maximum concordance with RNA binding preferences.

**Figure 1:**
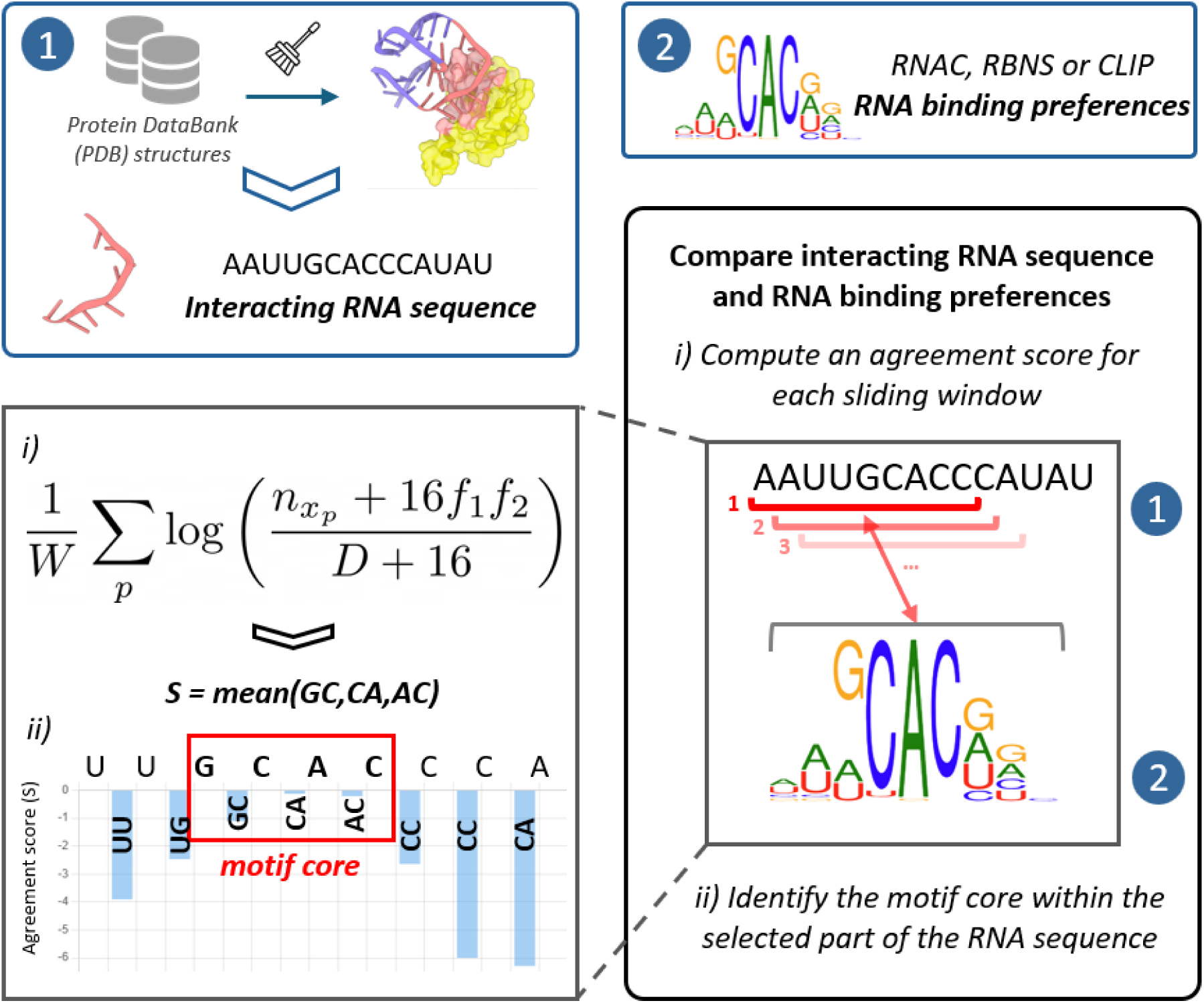
Workflow for scoring the agreement between structurally determined protein-binding RNA sequences and omics-based RNA binding preferences for that same protein. Information about protein-binding RNA regions was recovered from structural data (interacting RNA sequence) and omics experimental data (binding preferences represented as a logo). The RNA sequence interacting with the protein in the experimental structure was compared with the binding preferences (frequency matrices, see main text and Methods) from omics experiments by computing an agreement score at every position using a sliding window strategy (see Methods). In the agreement scoring formula (see Me, at a given dinucleotide position p, n_x:p_ represents the number of observed dinucleotides x in the RNA binding frequency alignment and D represents the total number of dinucleotides at this position (alignment depth). A pseudocount of 16 is added to manage dinucleotides with few occurrences. f1 and f2 represent normalized frequencies of the first and second nucleotides in x in the frequency matrix. W is the total number of dinucleotide positions. The best scoring sliding window was retained and the four highest-scoring consecutive nucleotides were assigned as the “motif core” for this interface.

The width of the frequency matrix depends on the number of columns in the 5-mer alignment, which makes it more difficult to compare agreement score values between proteins. Additionally, we saw that the edges of 5-mer alignments contain more gaps than the center of the alignment. Thus, a “motif core” was defined as the three consecutive dinucleotides (i.e. four consecutive nucleotides) yielding the maximal agreement score, assumed to represent the central region, key for the interaction with the protein.

The vast majority of the 294 PRI complexes exhibited near-zero agreement scores for RNAC and RBNS (median of -0.6 and -0.9 respectively) and lower scores for CLIP (median of -3.6) (Fig 2A). These results show that CLIP-identified motifs are less consistent with structural data, in agreement with the use of CLIP data only as a complement to RNAC and RBNS in our dataset. To filter relevant motifs based on agreement score values, we established a relevancy threshold (T) as the difference between the mean (μ) and the standard deviation (σ) of the scores (T=μ−σ), calculated from the combined RNAC and RBNS *in vitro* datasets to increase the statistical power of the threshold definition. We obtained a T value of -2.25 and removed a minority of RNAC and RBNS cases with agreement values below this threshold, resulting in a filtered dataset of 159/181 and 169/220 protein-RNA interfaces with RNAC and RBNS data, respectively. Using the same threshold, only 21/94 interfaces remain with CLIP data, covering 4 proteins, which represents insufficient data for further statistical analysis. Additionally, interfaces with CLIP values correspond partly to the less well-resolved structures in the dataset (Fig 2B). As a consequence, we did not consider CLIP data for further analysis.

**Figure 2:**
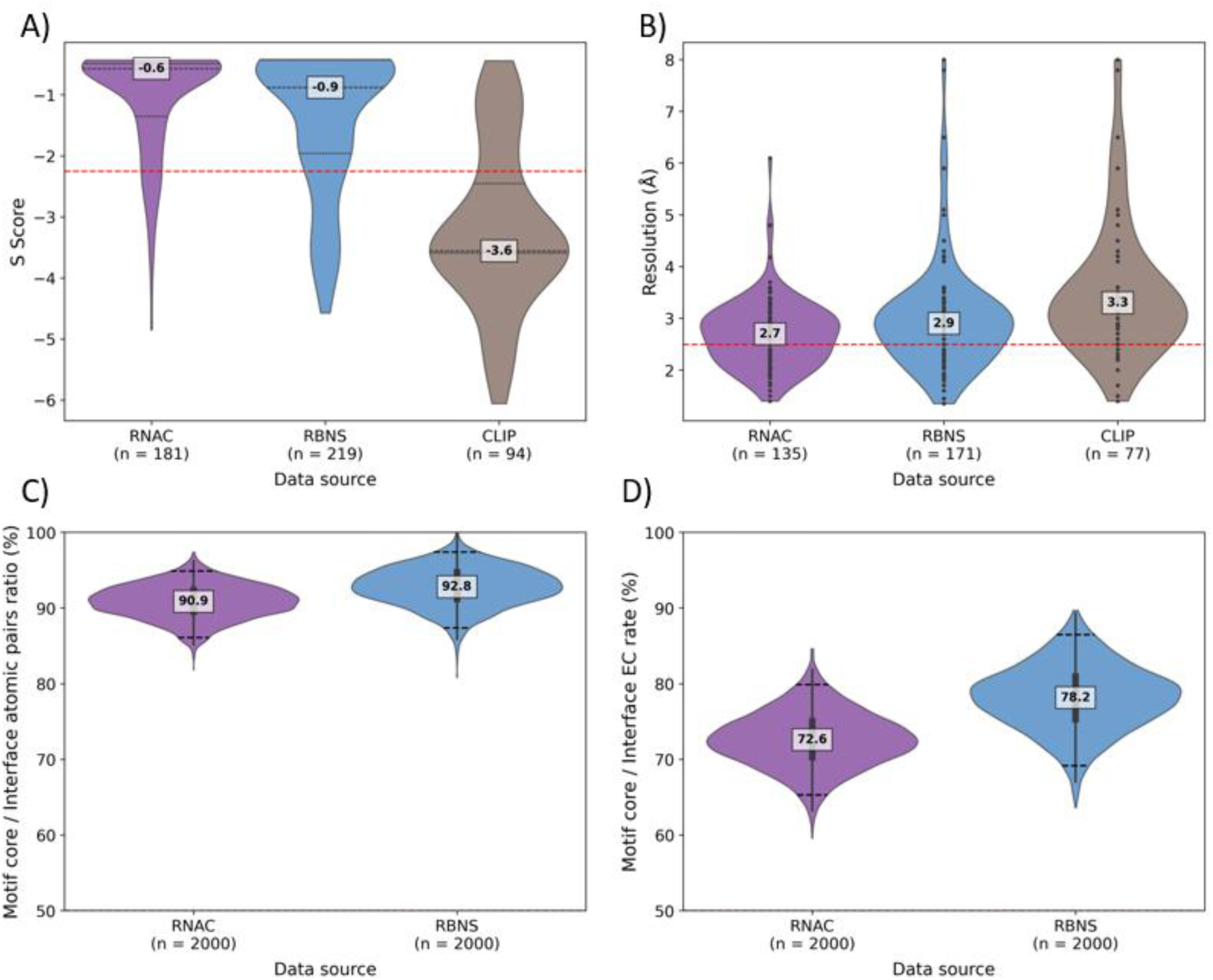
Agreement scoring and protein-RNA interaction dataset statistics. **A)** Violin plot with the agreement scoring results for motif cores of the 294 protein-RNA complexes including 181 with RNAC data, 219 with RBNS data and 94 with CLIP data. The dashed red line represents the threshold value T of -2.25. **B)** Violin plot of the structural resolution, when available, for subsets of interfaces with RNAC, RBNS and CLIP data. Structures with no available resolution value were determined by NMR spectroscopy. The dashed red line represents a standard resolution quality threshold of 2.5 Å. **C)** Violin plot of the distribution of atomic pairs (AP) ratio bootstrap values. The AP ratio is defined as the percentage of cases in which the average number of atomic contact pairs is higher in the motif core compared to the rest of the interface. Bootstrap values were obtained by drawing 2000 times with replacement from respectively 159 (RNAC) and 169 (RBNS) interfaces passing the agreement score threshold. **D)** Similarly to panel C, violin plot of the distribution of evolutionary conservation (EC) ratio bootstrap values. The EC ratio is defined as the percentage of cases (as in panel C, among interfaces passing the score threshold) in which amino acids in contact with the motif core are more conserved on average than amino acids of the rest of the interface.

To assess the robustness and structural relevance of the defined motif core, we investigated two key structural metrics: atomic pair (AP) contacts and evolutionary conservation (EC) of interacting residues (see Methods). We hypothesized that nucleotides belonging to the motif core should exhibit a higher density of contacts compared to the rest of the protein-RNA interface and that they should interact with more conserved amino acids, as these residues are likely under selection pressure to maintain the binding interface over evolutionary time.

The AP contact metric quantifies the average per-nucleotide number of atomic contacts within 5 Å of the protein chain in the 3D structure. This metric is highly correlated with solvent accessibility and provides insight into interface “anchors”, i.e. residues most tightly connected across the interface. On average, motif cores formed more atomic pairs than non-core nucleotides in 90.9% and 92.8% of the 159 (RNAC) and 169 (RBNS) cases, respectively (Fig 2C), with 95% confidence intervals from the bootstrap analysis (see Methods) of [86.1, 94.9] for RNAC and [87.4, 97.4] for RBNS. These results suggest that the motif core identified by the agreement score generally corresponds to the most structurally anchoring part of the RNA. Reduced AP contacts in some motif cores may result from different binding regions across structures, crystal packing artifacts, extended or dispersed interfaces. Atomic-level contacts are generally considered reliable for X-ray and cryoEM structures below 2.5 Å resolution; such well-resolved structures represent 32% and 30% of the RNAC and RBNS datasets, respectively (Fig. S2), while 24.5% of the 294 interactions were determined by NMR.

EC analysis similarly supported the structural relevance of motif cores. Protein residues interacting with the motif core were more conserved than those interacting with the rest of the interface nucleotides in 72.6% (95% CI [65.3, 79.9]) and 78.2% (95% CI [69.1, 86.5]) of RNAC and RBNS cases, respectively (Fig. 2D). Deviations may reflect the strict definition of motif cores, which can exclude interactions involving larger, dispersed, multi-site, or multi-stranded interfaces. As expected, interacting residues were also more conserved than general surface residues (93.1% and 95.6% for RNAC and RBNS, respectively, Fig S3).

To facilitate data visualization and integration, we developed a web interface (see Methods) providing access to all results for the 60 proteins and 294 structural interfaces analyzed in this study: https://bioi2.i2bc.paris-saclay.fr/rnaprotci/. For each protein, this user-friendly platform enables visualization of PDB structures, structural alignments, protein surface properties (solvent accessibility, electrostatics, evolutionary conservation), agreement scores, logos derived from frequency matrices, and identified motif cores. Crucially, it integrates these data with structural representations, allowing for interactive exploration of protein-RNA interactions. Users can search for proteins of interest using PDB identifiers, UniProtKB accession numbers, or keywords contained in the protein description.

### 2. Assessing protein-RNA interaction specificity in AlphaFold3 predictions

As we gathered protein–RNA complexes for which both high-resolution structures and experimental binding preferences are available, this dataset provides a valuable opportunity to evaluate the performance and specificity of AF3 predictions. In the original AF3 study, performance on protein–nucleic acid complexes was assessed on a set of 25 cases, with an estimated success rate of approximately 40%. Here, we extend this evaluation to a larger dataset, albeit unfiltered for redundancy with the AF3 training dataset.

Some PDB structures contain multiple copies of the same protein-RNA pair but with different interaction sites. Thus, from the 294 interfaces in the previously described dataset, we constructed a dataset of 280 unique protein-RNA pairs, on which AF3 was run to assess whether it could reproduce the experimentally observed interfaces. We found that all of these 280 protein-RNA pairs were redundant with the AF3 training data. 240 protein-RNA interactions were directly represented in the AF3 training set. The remaining 36 were not, but the proteins in these interfaces shared high structural similarity with proteins belonging to protein-RNA complexes released before the 30 September 2021 cutoff (see Methods).

We generated AF3 models (see Methods) for the 280 protein-RNA interfaces using protein sequences from their corresponding UniProtKB entries and RNA sequences derived from the associated experimental structures (“PDB-RNA” input). To systematically assess the ability of AF3 predictions to reflect experimentally defined specificity, we designed alternative RNA inputs for each complex. As a non-specific control, we used poly-U sequences matching the length of the PDB-derived RNA (“PolyU” input). In addition, we constructed semi-specific RNA sequences based on binding preferences derived from RNAC and RBNS experiments (“RNAC” and “RBNS” inputs, respectively). For RNAC and/or RBNS, a binding consensus was obtained from 5-mer enrichment analyses and embedded within flanking regions of 10 random nucleotides on each side to represent longer RNA contexts. For each input condition, we generated 5 AF3 models and retained the top-ranked model, as evaluated by the AF3 ranking score.

The median backbone root-mean-square deviation (RMSD) for the protein components of the complexes was 1.79 Å for AF3 top-ranked models, which reflects a high degree of structural accuracy for most predicted proteins, regardless of the input RNA sequence (Fig 3A). In contrast, the RNA predictions were markedly less accurate, with a median backbone RMSD of 5.1 Å and a large standard deviation (Fig 3B). These results highlight that the proteins in our dataset are folded into well defined domains, while the RNA may be more flexible. While statistically significant, the RNA RMSD difference between different input conditions (PDB-RNA, PolyU, RNAC, RBNS) is quite low. We then focused on the interface and analyzed the fraction of native contacts (fnat, see Methods) in AF3 models compared to the experimental structure (Fig 3C). The PDB-RNA models reached a median fnat of 0.56, significantly higher than the PolyU, RBNS and RNAC models.

**Figure 3:**
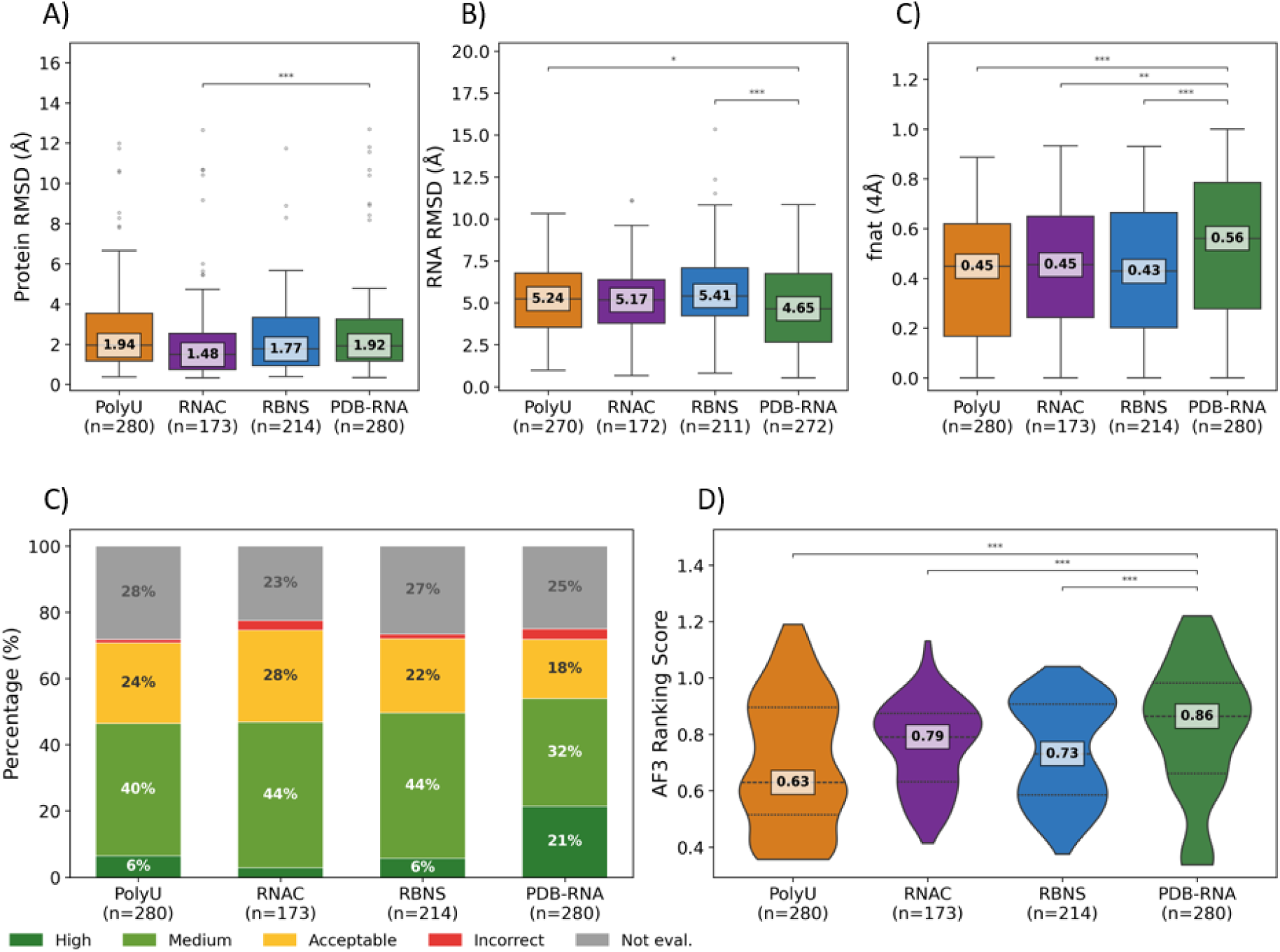
Evaluation of AlphaFold 3 specificity on protein-RNA interactions in our dataset compared to reference structures. **A)** Boxplots showing the root mean square deviation (RMSD) of the protein backbone for the 280 top-ranked AF3 models obtained using four RNA input types: PolyU, RNAC, RBNS and PDB-RNA. The central line and boxed value represent the median. **B)** Same as panel A, for the RNA RMSD. **C)** Boxplots showing the fraction of native contacts (fnat) for the 280 top-ranked AF3 models obtained using four RNA input types. The central line and boxed value represent the median. **D)** Systematic evaluation of AF3 models for all conditions using CAPRI metrics, classifying predictions into High quality (dark green), Medium (light green), Acceptable (orange) or Incorrect (yellow). Models classified as Not Evaluated (grey) correspond to cases where fewer than 5 protein or 5 nucleic acid residues were identified within the interface contact threshold, making interface quality metrics undefined. **E)** AF3 ranking scores for top-ranked models in each of the four conditions. The central line and boxed value represent the median. Pairwise statistical comparisons against PDB-RNA were performed using two-sided Mann-Whitney U tests (* p < 0.05, ** p < 0.01, *** p < 0.001).

We further applied the CAPRI (Critical Assessment of PRediction of Interactions) success criteria (Fig 3D), with protein-protein thresholds as a proxy for protein-nucleic acid ones [25] (see Methods). For all four input RNA sequences, a notable fraction of 23% to 28% of predictions could not be evaluated, due to the structural alignment covering fewer than 5 interface residues on either the protein or the RNA side, which is insufficient for reliable CAPRI metrics computation, and effectively means that AF3 does not position the interface correctly. Depending on the RNA input condition, 1 to 3% of predicted interfaces proved to be Incorrect (I). Including Acceptable or better models, the predictive performance of AF3 for protein-RNA complexes was 70% to 75% depending on the RNA input condition, showing that the AF3 success rates were very similar regardless of the RNA sequence we used for the prediction. However, PDB-RNA based models got the highest proportion of High quality models (21%, vs. 3% to 6% for other conditions), demonstrating that the ability of AF3 to predict precise interface contacts seems to depend on the use of the exact RNA sequence seen during training. The average AF3 ranking score, reflecting the internal AF3 confidence, is also significantly higher in the PDB-RNA condition (Fig 3E). Similar trends held when the SNRPA protein (involved in 75/276 modeled interfaces) was removed from the analyzed dataset (Fig S4).

To assess whether the observed trends are influenced by interactions with omics-based binding preferences poorly matched to the structural RNA sequence, we repeated the analysis after applying the agreement score filter (S > −2.25, see Section 1), retaining 236 interactions for which at least one experimental technique showed significant concordance between the binding preferences and the interface structure (Fig S5). After filtering, AF3 ranking scores for RBNS increased moderately (from 0.73 to 0.80), which could be expected, while RNAC remained stable (0.79) and the PDB-RNA advantage in fnat and CAPRI classification persisted. This indicates that the advantage of PDB-RNA inputs is not driven by cases with poorly matched binding preferences. The tendencies we observed may reflect either the sensitivity of AF3 to the exact RNA sequence encountered during training, which might point to signs of memorization rather than sequence level generalization, or a genuine importance of the nucleotidic context around the motif core to define the binding specificity.

### 3. Detailed case studies highlight some limitations of the agreement score and AlphaFold3 predictions

#### 3.1 SNRPA: Agreement scoring identifies the interaction motif core, while AF3 shows limited specificity and signs of memorization

We focused on the most represented protein in our dataset, the U1 small nuclear ribonucleoprotein A (SNRPA). This protein contains two RNA-binding domains (RRM1 and RRM2), and is known to preferentially interact with “UGCAC” RNA motifs. 93 structural complexes involving SNRPA are present in the PDB, all containing the RRM1 domain, with high protein structural similarity (RMSD of the protein backbone relative to reference structure 1AUD: 1.0 to 2.31 Å) and sequence similarity (sequence identity with human SNRPA: 87.5% to 100%). 75 out of the 93 complex structures are kept for this study after interface size filtering (See Methods).

The RNA interacting motif identified from RNA binding frequency data highlights a strong “GCAC” specificity for both RNAC and RBNS experimental methods (Fig 4A,B). The agreement scoring workflow successfully identified this specific motif in 68 protein-RNA complexes sharing high structural similarity (with iRMSD between 1.0 and 4.2 Å) and high agreement scoring values: -0.49 (RNAC) and -0.42 (RBNS). Moreover, both AP and EC are enriched in those identified motif cores, except in the 3L3C interface, due to minor distance variation in the structure.

**Figure 4:**
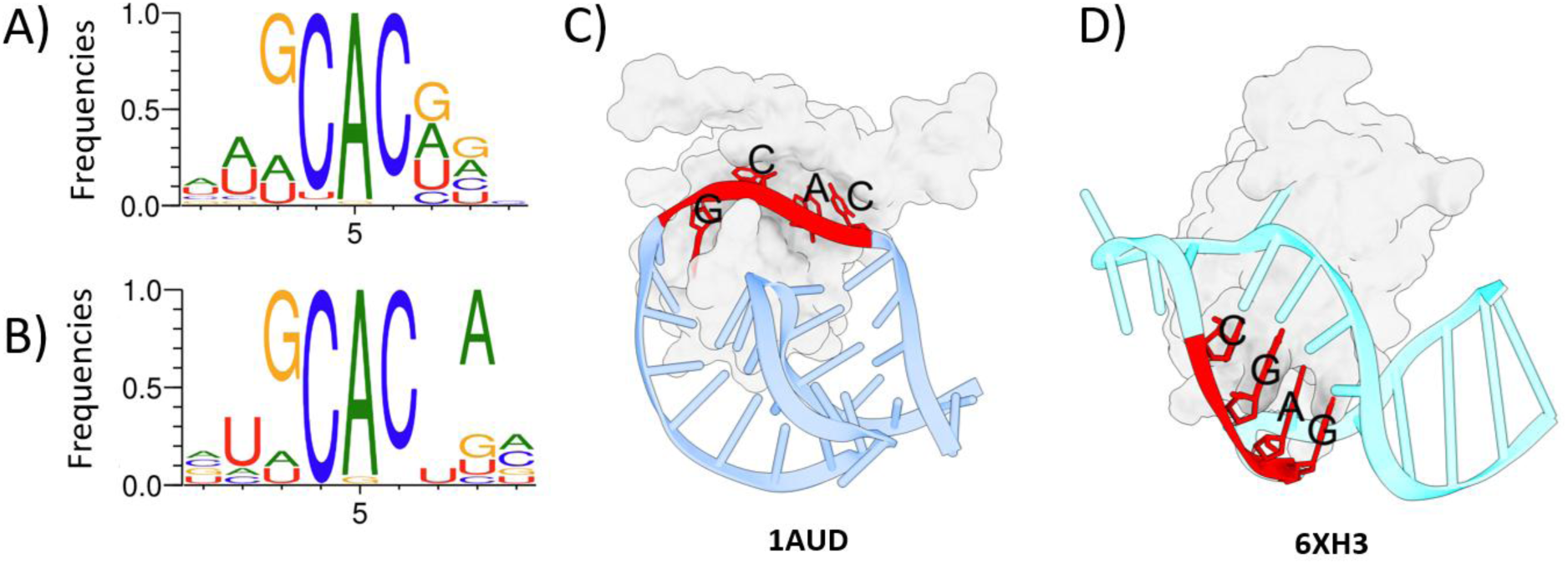
Overview of SNRPA RNA binding specificity. **A)** RNAC binding profile obtained from the alignment of 39 5-mers. **B)** Same for RBNS, from 18 5-mers. **C)** Structural overview of the 1AUD protein-RNA interface. The motif core is represented in red. **D)** Structural overview of the 6XH3 protein-RNA interface. The motif core is represented in red.

The remaining 7 structures harbor a wide score range (from -0.6 to -2.8); the more the identified motif core differs from “GCAC”, the lower the score (Table S1). Analysing these cases in more detail enhanced our understanding of the agreement score’s behavior. First, four almost identical PDB structures (6XH0, 6XH1, 6XH2 and 6XH3, iRMSD between 0.3 and 0.8 Å) contain a different RNA-binding site from other SNRPA structures, synthetically modified in order to ease crystallization [26]. The RNA sequence is also different as it corresponds to variants of the HIV-1 Trans Activation Response element (TAR), leading to structural differences compared to reference structure 1AUD (Fig 4C,D). In these four structures, the agreement scoring workflow identified “GAGC” and “AGCU” motif cores associated with scores of -2.44 and -2.79, from RNAC and RBNS data respectively, both lower than the -2.25 threshold. These cases are thus successfully identified as true negatives by our agreement scoring workflow. Moreover, they form fewer AP contacts with motif cores than with the rest of the structural interface and motif core-interacting amino acids are less evolutionarily conserved. On the other hand, for the 4YB1 PDB structure, the workflow identified “CACA”/“NCAC” motif cores with good scores (−0.68/−0.76 in RNAC/RBNS), despite an incorrect biological interface. Structural inspection revealed a symmetry-related interface mediated by a “GCAC” motif, suggesting that the biological assembly of 4YB1 was incorrectly assigned in the PDB.

We assessed AF3 performance for SNRPA complexes using input sequences representative of the two experimentally observed binding modes, from PDB structures 1AUD and 6XH3. AF3 reproduced the protein structures reasonably well, with RMSDs of 2.1 Å and 1.3 Å for the top-ranked 1AUD- and 6XH3-based models, respectively, whereas RNA RMSDs were higher (5.1 Å and 2.6 Å). The difference between RMSD values for 1AUD and 6XH3 motivated a more detailed assessment of the predicted interfaces.

For 1AUD, AF3 consistently reproduced the experimental binding mode. PDB-RNA-based models showed especially high ranking scores (0.97–0.99) and recovered the strong “GCAC” motif as the main anchored RNA region. PolyU-, RNAC- and RBNS-derived sequences led to lower ranking scores, but still largely reproduced the 1AUD-like interface and recovered the “GCAC” region in RNAC- and RBNS-based models (Fig 5A, Table S2). iRMSD values remained comparable across conditions (1.34–1.80 Å), indicating that this binding mode was robustly predicted despite changes in the input RNA sequence.

**Figure 5:**
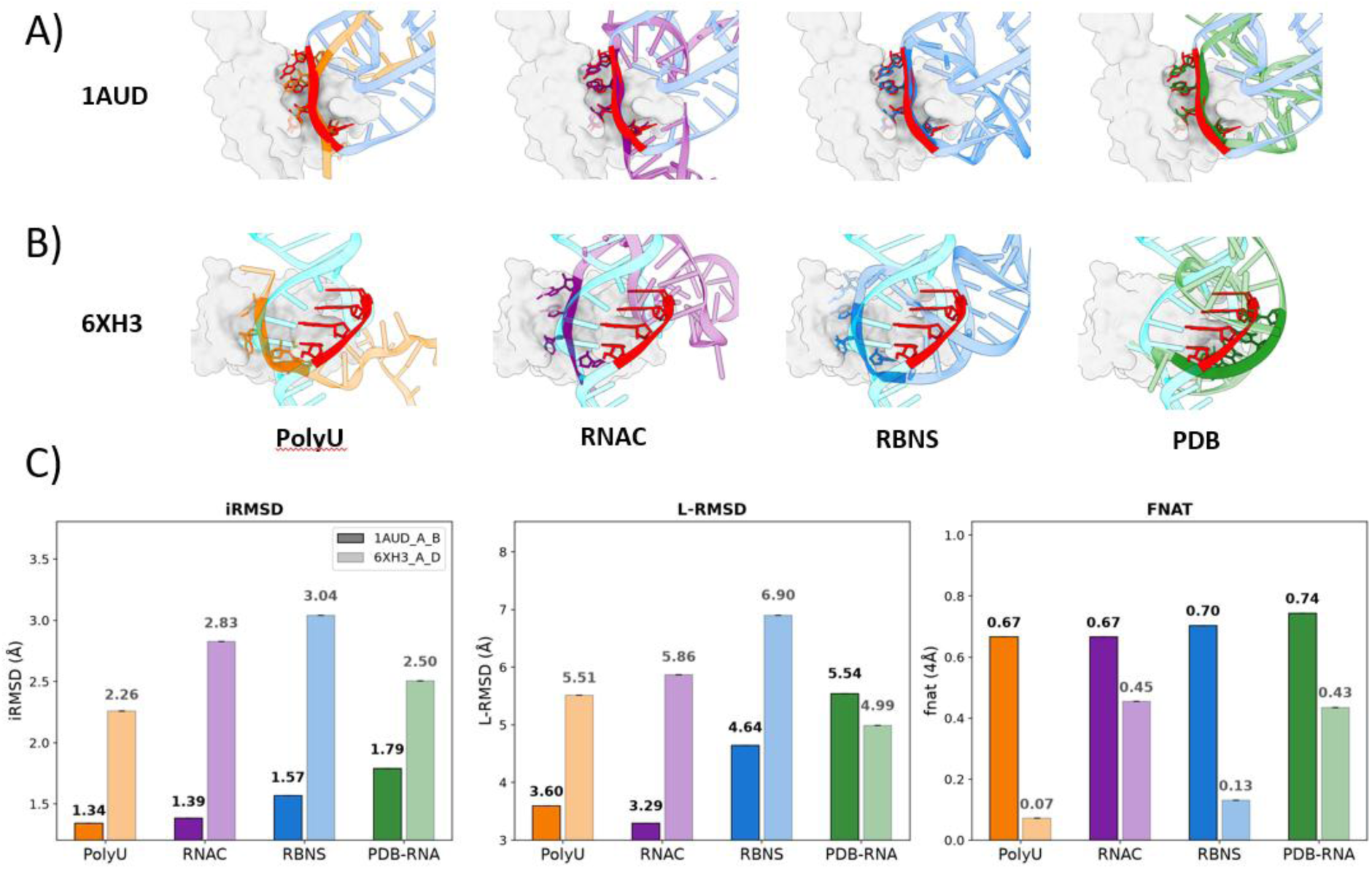
AlphaFold3 specificity analysis for SNRPA protein-RNA interaction prediction. **A)** Zoom on the 1AUD PDB protein-RNA interface (motif core colored in red, rest of the RNA in light blue) superimposed with AF3 generated models using different RNA sequence (PolyU RNA in orange, RBNS RNA in marine blue, RNAC RNA in purple, PDB RNA in green). **B)** Same as A) using 6XH3 as reference. **C)** iRMSD, L-RMSD and fnat values of top-ranked AF3 model for each RNA input condition, compared to 1AUD and 6XH3 references.

In contrast, AF3 showed limited ability to recover the alternative 6XH3 binding mode. The top-ranked PDB-RNA-based 6XH3 model received a lower ranking score (0.76) and adopted an intermediate conformation rather than reproducing the experimental 6XH3 interface (2.5 Å iRMSD compared to 6XH3). PolyU-, RNAC- and RBNS-based models instead tended to resemble the 1AUD-like interface (Fig 5B). This preference is also reflected in RNA contact prediction: the top-ranked 1AUD model reached 83% precision and 100% coverage, compared with 50% precision and 62% coverage for the 6XH3 model, while protein-side precision and coverage remained around 80% for both (Fig S6). Interestingly, AF3 could still recover several correct interacting residues on the protein side in 6XH3-based models, suggesting that the main limitation lies in reproducing the RNA-binding mode rather than the protein interface itself.

We further validated our interface analysis with a CAPRI-like evaluation of the models, with markedly poorer iRMSD, L-RMSD and fnat values for 6XH3 than for 1AUD across conditions (Fig 5C). All top-ranked 1AUD models reached the Medium CAPRI category, whereas only the PDB-RNA-based 6XH3 model was classified as Medium and the remaining models as Acceptable. Overall, AF3 preferentially converged toward the 1AUD-like binding mode, showing limited sensitivity to the input RNA sequence and reduced ability to recover the alternative 6XH3 interface. This suggests a potential bias toward a favored binding mode, possibly reflecting its stronger representation in the training data.

#### 3.2 LIN28A: Alternative interaction interfaces remain challenging to identify with the agreement score and AlphaFold3

LIN28A illustrates the challenge of analyzing proteins with multiple RNA-binding sites. Two structures, 2LI8 and 5UDZ, represent distinct RNA-binding interfaces of LIN28A, with the protein Zinc Knuckle Domain (ZKD) for 2LI8 and the Cold Shock Domain (CSD) for the 5UDZ biological assembly (Fig 6A). RNAC experiments show that LIN28A recognizes RNA sequences including the “GGAG” motif (Fig 6B), consistently with the existing literature [27, 28]. In 2LI8, our scoring workflow correctly identifies “GGAG” as the motif core (score = −0.5) and this motif is central in the interface structure (Fig 6C). In contrast, for 5UDZ, the workflow identifies an “NGGU” motif (score = −1.7) that does not correspond to the most structurally anchored RNA region (Fig 6D). This RNA region is less conserved and makes fewer AP contacts than the interface in 2LI8. This discrepancy likely results from the coexistence of different RNA-binding signals for each interface, which might be merged in the experimental binding profile (Table S3).

**Figure 6:**
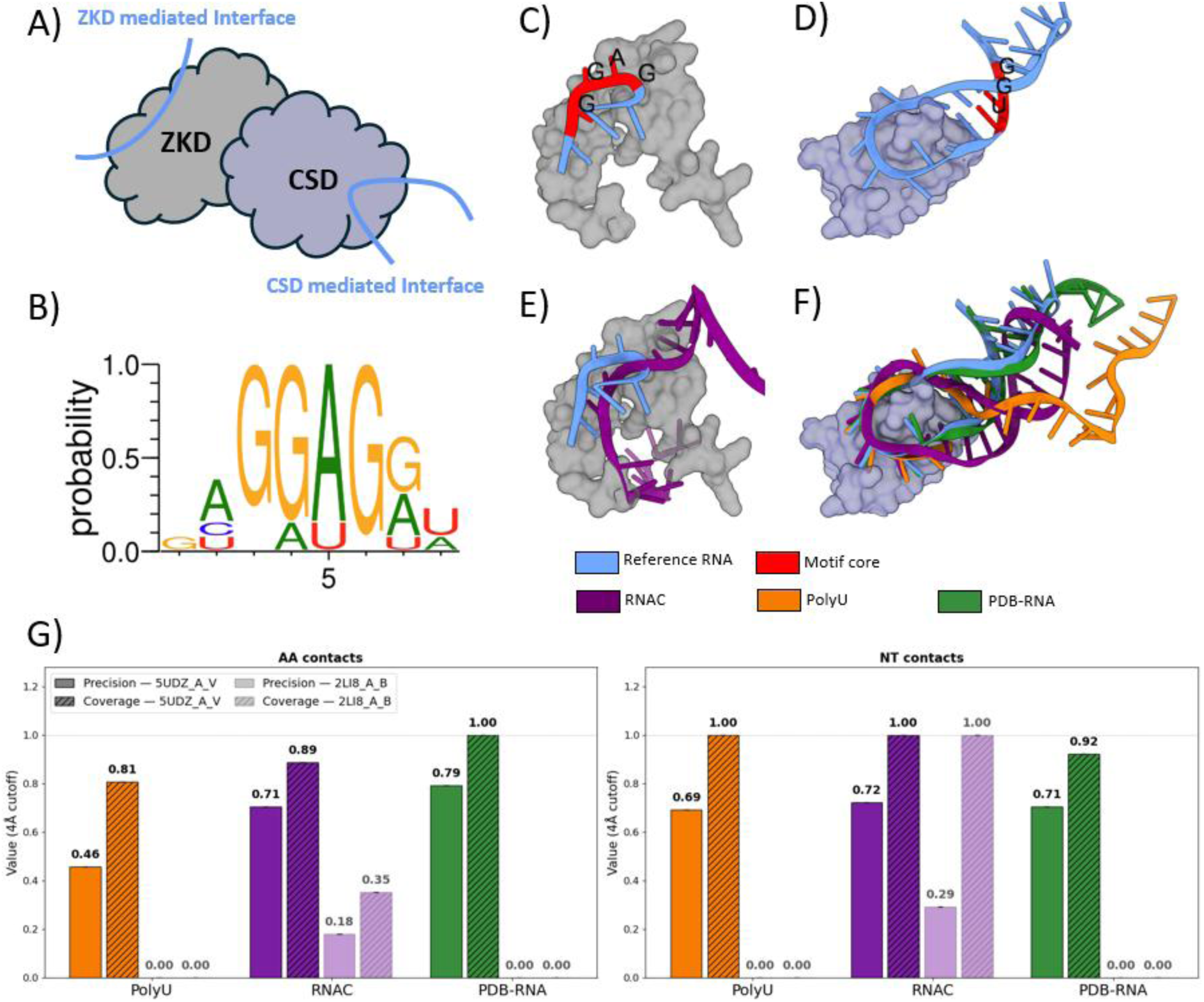
LIN28A contains two RNA binding domains. A) Schematic representation of the LIN28A structure and RNA-binding sites. B) RNAC binding profile obtained after 5-mer alignment. C) Structure of the LIN28A protein in complex with RNA at the Zinc Knuckle Domain (ZKD) binding site (PDB ID: 2LI8). The identified motif core is colored in red. D) Structure of the LIN28A protein in complex with RNA at the Cold Shock Domain (CSD) binding site (PDB ID: 5UDZ). The identified motif core is colored in red. E) LIN28A 2LI8-based AF3 model superimposed with reference RNA (from the PDB structure). Only the RNAC model is shown as RNAs from other conditions were predicted to interact with the other domain of the protein. F) LIN28A 5UDZ-based AF3 models with the three RNA input conditions, superimposed with reference RNA (from the PDB structure). G) Precision and coverage metrics for amino acid and nucleotide contacts in top-ranked AlphaFold3 models, over three RNA input conditions.

AF3 showed a similar discrepancy. Although AF3 ranking scores are higher for PDB-RNA-based models (Table S4), the difference in prediction quality is less pronounced. For 5UDZ, AF3 consistently reproduced the CSD-mediated interaction, with >80% coverage, independently of the RNA sequence. Conversely, AF3 generally failed to reproduce the ZKD-mediated interaction observed in 2LI8. The RNAC-derived sequence was an exception, with 100% coverage but only 29% precision, indicating that the RNA was positioned at the correct surface without reproducing the specific interaction (Fig 6E–G). However, the AF3 ranking score was high (Table S4). Together, these results highlight the difficulty of recovering RNA-binding specificity when a protein contains multiple binding interfaces.

## DISCUSSION

The present study introduces a scoring workflow to assess the agreement between experimental RNA binding preferences and structural information, in view of providing guidelines for analyzing protein–RNA interaction specificity. By uniquely combining independent data sources, the score provides a quantitative framework for identifying the RNA nucleotides most likely to mediate protein recognition, which we designate as “motif core”. This makes it especially interesting for the qualification of upcoming data in view of enriching existing benchmarks.

The observed enrichment of atomic contacts and evolutionary conservation within the identified motif cores further supports the ability of our approach to reliably distinguish biologically relevant interaction regions. Detailed case studies of SNRPA and LIN28A interactions with RNA illustrate the robustness of our workflow, as most cases demonstrate its value to identify a relevant motif core. The LIN28A analysis also showcases some difficulties linked to the presence of multiple protein-RNA binding sites. Beyond the scope of this study, results might be improved by performing protein-specific selection of relevant 5-mers from RNAC or RBNS, in order to build binding profiles better representing the true binding preferences.

In a context where database and web-based tools dedicated to protein-nucleic acid interactions are not as developed as those focused on protein-protein interactions [29], we provide access, through a web interface including interactive visualization of 3D interface structures, to detailed agreement scoring results on the dataset from the present study: https://bioi2.i2bc.paris-saclay.fr/rnaprotci/ This user-friendly interface can be browsed by type of available binding preferences, by PDB identifier or through a broader keyword search.

We leveraged our dataset combining structural and binding preference data to analyze the ability of AlphaFold3 to accurately predict protein-RNA interaction specificity. In most cases, AF3 models reproduce the experimental interface, showing great backbone recovery when compared to the reference structure. Because our dataset contained structures that the model encountered during its training phase, these high-performing results must be interpreted carefully with respect to training-set overlap and do not constitute an independent evaluation of AlphaFold3 performance on de novo protein-RNA prediction. However, thanks to the unique combination of structural and binding preference data, we showed that the performance of AF3 is almost equally good when given as input the RNA sequence from the experimental structure, a sequence containing the binding motif from experimental binding preferences, or a non-specific polyU sequence, demonstrating that AF3 struggles with RNA sequence-level specificity. Those tendencies are very similar whether or not the benchmark dataset was filtered to retain only cases with good agreement between the experimental binding preferences and the binding region in the PDB structure.

This is in line with the evaluation of protein-RNA prediction in CASP16, where the prediction accuracy for those complexes was shown to depend on template availability [30], as well as two other recent studies of protein-RNA interface prediction by AlphaFold3. Ludaic & Elofsson showed that RNA and protein-RNA structure prediction are still very challenging for both AlphaFold3 and other deep-learning methods, and that these methods tend to perform much better when the target is similar to structures seen by the method during training [31]. This indicates that the model’s success relies on recognizing previously observed structural patterns, rather than generalizing to novel folds and interfaces. In another recent paper [32], Follonier et al. confirmed that AF3 performance depends strongly on the presence of related complexes in the training set; they additionally showed that AF3 often fails to predict the correct specificity, in particular for protein-RNA interactions where the confidence metrics cannot be trusted to separate genuine from random target RNA sequences. These findings are very consistent with ours.

One remaining limitation of the current study lies in the availability of suitable experimental data. Our benchmark for the agreement scoring workflow and AlphaFold3 prediction specificity is currently restricted to human proteins for which both RNAcompete or RNA Bind-n-Seq data and experimentally determined protein–RNA structures are available, resulting in a relatively small dataset. RNAcompete and RNA-Bind-N-Seq data remain scattered across the literature. Extending our approach to integrate *in vivo* RNA binding preferences would improve its ability to reflect biological implications of protein-RNA interactions, but remains challenging because of scarce and lower-quality data. Our agreement scoring workflow will facilitate the qualification of new experimental binding data in order to leverage future generalization and enrichment of RNA-binding repositories.

Our study opens perspectives regarding possible use of orthogonal sources of information to improve protein-RNA structure prediction. One interesting direction would be to test whether omics-derived motifs can be used as constraints to guide modeling. A valuable tool to test this might be Protenix, one of the few methods currently supporting user-defined distance constraints [33]. More generally, findings from the present study might be extended to improve the training and performance of protein-RNA structure predictors by building enriched MSAs incorporating RNA binding preference information, and to develop prediction models fine-tuned for specificity. The latter idea, although previously expressed by others [9, 34], remains to be implemented in practice as it still suffers from several of the aforementioned challenges, in particular the scarcity of structural and omics-derived information.

## METHODS

### 1. RNA binding frequencies pre-processing

RNA binding frequencies were derived from pre-processed RNA compete (RNAC), RNA Bind-n-Seq (RBNS) and Cross-linking Immunoprecipitation (CLIP) data [35], represented as lists of RNA 5-mer sequences with associated 5-mer enrichment scores for 381 protein identifiers. 35 proteins with only CLIP data, lacking both RNAC and RBNS data, were discarded.

To select a list of most relevant 5-mers, score distributions were smoothed using “np.convolve” (NumPy 1.26.4 [36], default parameters) before identifying a strict inflection point via “np.diff”. 5-mers with scores exceeding this threshold were aligned using MAFFT [37] (default parameters, --auto) and this alignment was designated as the seed alignment. Subsequently, a less stringent inflection point was determined using the “KneeLocator” function from the kneed package [38], defining a list of additional 5-mers that were then aligned to the fixed seed alignment using MAFFT with the “--reorder” and “--addfragment” parameters. A protein-specific frequency matrix, specifying the proportion of each possible nucleotide at each position, was derived from each resulting alignment, including gaps, so that the matrix size depends on the alignment length. WebLogo [39] was used to build a visual representation of each frequency matrix. Another protein-specific, dinucleotide-based, frequency matrix was computed from each alignment for later dinucleotide-based scoring.

### 2. Pre-processing of structural data and RNA binding preferences pre-processing

Protein-RNA complexes were extracted from the Protein Data Bank (PDB) [6, 7] dated 06-11-2024, leading to 5704 structure identifiers and, after biological assembly reconstruction, 173,679 interacting pairs of protein and RNA chains (with protein-RNA interaction defined by heavy atom distance lower than 5 Å). RNA binding frequencies and protein-RNA interaction structures were mapped together by directly extracting UniProtKB [40] accession numbers from annotated PDB entries: we obtained 494 PDB entries containing a protein with RNA binding frequency data from RNAC and/or RBNS. In parallel, for all PDB-derived interfaces, a BLAST search [41] was performed using the protein sequence as a query, down to a 50% identity threshold to identify closely homologous proteins with RNA-binding frequency data; this additional step recovered 60 additional PDB entries. Protein-RNA structural complexes with fewer than 5 nucleotides or 5 amino acids at the interface were excluded from the dataset. Additionally, proteins shorter than 50 amino acids were removed, leading to a dataset of 294 pairs of interacting protein and RNA chains with associated RNA binding preference data.

### 3. Agreement score

The RNA sequence for all nucleotides at less than 5 Å (heavy atom distance) from the protein in the three-dimensional PDB structure was extracted and referred to as the interacting RNA sequence. Gaps between interacting positions that are not consecutive in the RNA sequence are filled with Ns. The agreement score S (Fig S1a,b), inspired from a transcription factor binding site predictor using position weight matrices [24], was computed by aligning the dinucleotide-based frequency matrix (from RNA binding preferences) to subparts of the interacting RNA sequence using an iterative sliding window strategy (Fig 1). We used dinucleotide-based scoring to better capture binding preferences. Each element *L* is defined as the likelihood to observe dinucleotide *x* at position *p* in the interacting RNA sequence subpart, with respect to the dinucleotide-based frequency matrix and 5-mer alignment:

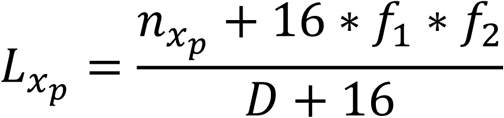

At a given dinucleotide position *p*, *n_x:p_* represents the number of observed dinucleotides *x* in the RNA binding frequency alignment and D represents the total number of dinucleotides at this position (alignment depth). A pseudocount of 16 is added to manage dinucleotides with few occurrences: *f1* and *f2* represent normalized frequencies of the first and second nucleotides in *x* in the frequency matrix, and there are 16 possible dinucleotides.

This score is applied to all dinucleotides (we defined W as the total number of dinucleotide positions) at all positions of the given sliding window in the RNA sequence and averaged to simplify comparisons between proteins:

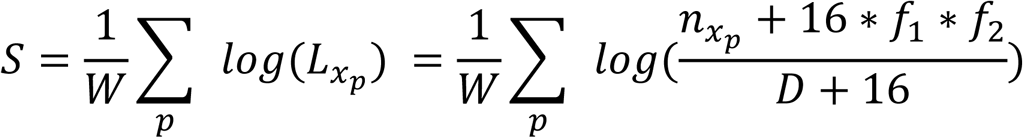

### 4. RNA binding frequencies and structural data agreement scoring workflow

The agreement scoring workflow begins with a scanning phase, using a sliding window approach. First, the dinucleotide-based frequency matrix (representing RNA binding preferences) is aligned to the start of the interacting RNA sequence, and the agreement score is calculated. The frequency matrix is then shifted by one nucleotide, and the score is recalculated. This iterative process continues until all possible positions of the frequency matrix across the interacting RNA sequence have been scored. The aligned subpart of the interacting RNA sequence with the highest score is selected, as it shares the best agreement with RNA binding preference data. Following the scanning phase, we analyze the contribution to the overall score of each dinucleotide in the best-scoring window. The three consecutive (overlapping) dinucleotides with the highest contributions are identified, and the four corresponding nucleotides are extracted to define the “motif core” of the interaction.

To handle alignment of frequency matrices and RNA subsequences at the edges of the RNA interacting sequence, a number of “N” characters equal to the ceiling of half the RNA sequence length is appended to both ends of the interacting RNA sequence. This allows for scoring the alignment of the initial positions of the interacting RNA sequence with the terminal positions of the frequency matrix, and vice versa. Similarly, if interacting RNA sequences contain positions that are non-contiguous in sequence, nucleotides that are missing because they do not interact with the protein are replaced with “N” characters. The scoring of “N” characters is performed using the mean frequency over all four bases at that specific position.

### 5. Data bootstrapping

Bootstrapping analysis was performed on the 178 and 199 filtered cases for RNAC and RBNS respectively. Subsequently, a bootstrap sample was generated by randomly selecting, with replacement. For each bootstrap sample, differences in evolutionary conservation and atomic pair (AP) contacts were calculated, with specific comparisons made between the motif core and the rest of the interface, and between the interface and the protein surface. This process was repeated for n = 2000 iterations. Finally, the average of the differences in evolutionary conservation and AP contacts was calculated across all 2000 iterations to obtain a representative value for each comparison.

### 6. Protein-RNA interaction features: protein evolutionary conservation and atomic pairs

For each protein, homologous sequences were retrieved using HHblits [42] against the Uniref30 version [43] dated February 2022. Redundancy was filtered with HHfilter [44, 45], using 25% minimum sequence identity and a diff parameter of 100, limiting the number of sequences while ensuring sequence diversity in the MSA. Evolutionary conservation was mapped from the MSA onto the structural models using conservation scores computed with Rate4Site [46]. Atomic pairs (AP) were obtained running x3DNA v2.4 [47] with default parameters on PDB structures.

### 7. Protein-RNA web interface

A webserver hosted at https://bioi2.i2bc.paris-saclay.fr/rnaprotci/ proposes a user-friendly interface to navigate over protein-RNA interaction data. Django v4.1.7 [48] and NGL viewer [49] were used for graphical interface development. Protein-RNA complexes are referred to as the four-letter code of the PDB and associated protein and RNA chain names (ex: 1AUD_A_B).

### 8. AlphaFold3 modeling of protein-RNA interactions

Protein-RNA interactions were modeled with local AlphaFold3 v3.0.0 built on our team server from the DeepMind github, with an A100 80 Gb GPU. Input protein sequences were extracted from their UniProtKB accession number, and RNA sequences from the structural PDB interface file. For models using a polyU RNA, we used the same length as the PDB RNA. For RNAC and RBNS based models, we extracted the consensus (most frequent nucleotide at each position) from the initial 5-mers alignment and embedded this consensus in a longer sequence by flanking it with two 10nt random sequences. MSAs were generated using the default parameters in AlphaFold3. Predictions were carried out using a single seed, without using templates, and produced 5 ranked models for each condition.

To assess the redundancy of our dataset with the AF3 training dataset, we extracted cases released after the 30-09-2021 cut-off and structurally compared them to the AF3 training set using Foldseek [50]. Structures were considered as never seen by AF3 during training when Foldseek was unable to find similar proteins interacting with RNA and having a “prob” score was under 0.5 and the e-value above 0.05.

### 9. Similarity between experimental and AlphaFold3-generated protein-RNA interactions

Experimental and modeled structures were aligned using GTcomplex (v1.0.9) [51] for structure-based residue-level correspondence. We computed fnat, iRMSD, and L-RMSD following CAPRI criteria for protein-protein interfaces [25]. The fraction of native contacts (fnat) was defined as the fraction of experimental amino acid-nucleotide pairs recovered in the prediction, using the gemmi library [52] with a 4Å distance threshold on all heavy atoms. The interface RMSD (iRMSD) was computed over protein and RNA residues within 8Å of their partner chain, using Cβ atoms for protein and C3’ for RNA. The ligand RMSD (L-RMSD) was computed using USalign (version 2024-11-08) with protein-only superposition at 5Å of the RNA chain, using Cα atoms. CAPRI categories were assigned as for protein-protein interfaces: High: fnat >= 0.5 and (L-RMSD <= 1.0 or iRMSD <= 1.0)). Medium: either 0.3 <= fnat < 0.5 and (L-RMSD <= 5.0 or iRMSD <= 2.0) or fnat >= 0.5 and L-RMSD > 1.0 and iRMS > 1.0. Acceptable: either 0.1 <= fnat < 0.3 and (L-RMSD <= 10.0 or iRMSD <= 4.0) or fnat >= 0.3 and L-RMSD > 5.0 and iRMSD > 2.0). Incorrect: fnat < 0.1 or L-RMSD > 10 and iRMSD > 4.0.

In addition, precision (fraction of predicted contacts found in the experimental structure) and coverage (fraction of reference contacts found in the prediction) were computed at both amino acid and nucleotide residue levels, using a 5Å threshold to define the interface, with Cα atoms for protein and C3’ for RNA.

### 10. Case studies

The reference sequence for SNRPA was obtained from the UniProt identifier P09012. For SNRPA, we arbitrarily chose 1AUD as the reference structure for the analysis, as it appropriately represents most SNRPA structures in association with RNA we found in the PDB. The reference sequence for LIN28A was obtained from the UniProt identifier Q9H9Z2.

Structural visualisation and analysis were done using ChimeraX v1.10.1 [53]. PyMol 3.2 [54] was also used for analysing the crystal packing interface of PDB ID 4YB1 with the symmetry mates function.

## ACKNOWLEDGMENTS

We thank Raphael Guerois for fruitful discussions and feedback on the manuscript. We thank Carla Martins for early exploration of the binding preference data and agreement scoring.

