## Supplementary Tables and Figures for "Bridging the gap between omics and structural data: A framework for interpreting protein-RNA interaction specificity"

**Short title: Protein-RNA binding specificity through omics, structure and AlphaFold3**

Yoann Fauconnet[1], Shuyun Deng[1], Rebecca Koundri[1], Martin Pagani[1], Chloé Quignot[1], Déborah Antonio[1], Jessica Andreani[1*]

[1]: Université Paris-Saclay, CEA, CNRS, Institute for Integrative Biology of the Cell (I2BC), 91198, Gif-sur-Yvette, France.

### Supplementary Figures

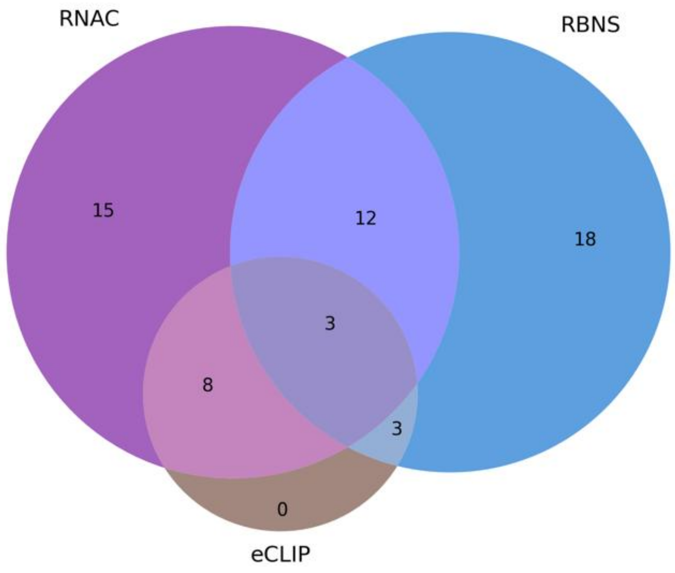


***Figure S1: Venn diagram representing the number of proteins with RNAC, RBNS and CLIP data among the 60 unique proteins present in our dataset of 294 structurally resolved protein-RNA interfaces.*** *RNAC stands for RNAcompete, RBNS for RNA-Bind-N-Seq and CLIP for Cross-Linking Immuno Precipitation assay.*

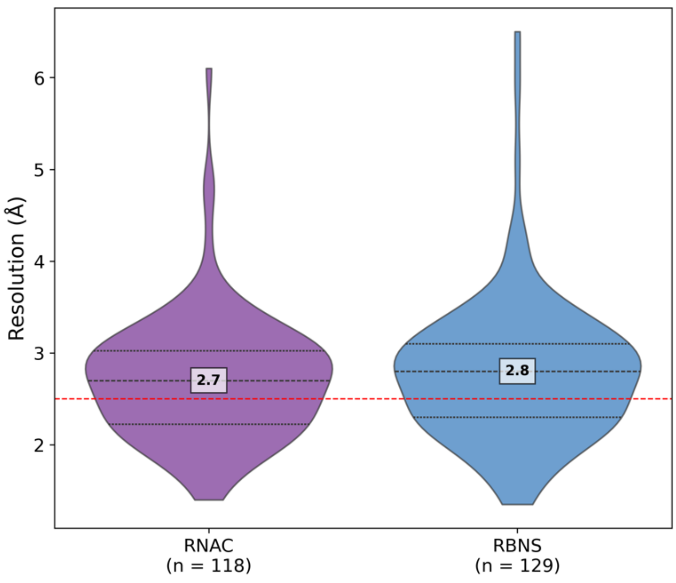


***Figure S2: Violin plot of the structure resolution of the dataset filtered with a S=-2.25 score threshold.***

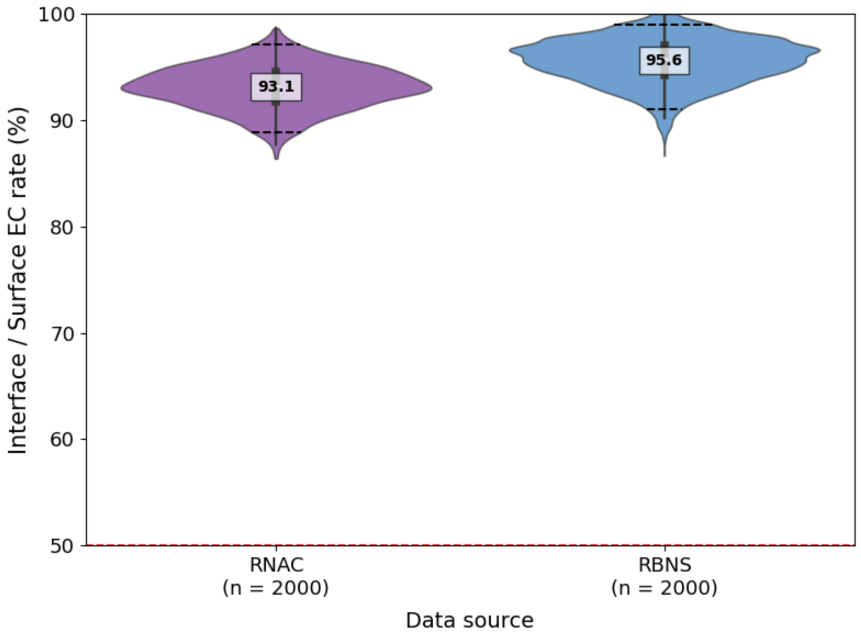


***Figure S3: Bootstrap analysis of protein interface versus surface Evolutionary Conservation analysis.*** *Violin plot of EC interface to surface ratio bootstrap results (N = 2000) for both RNAC and RBNS.*

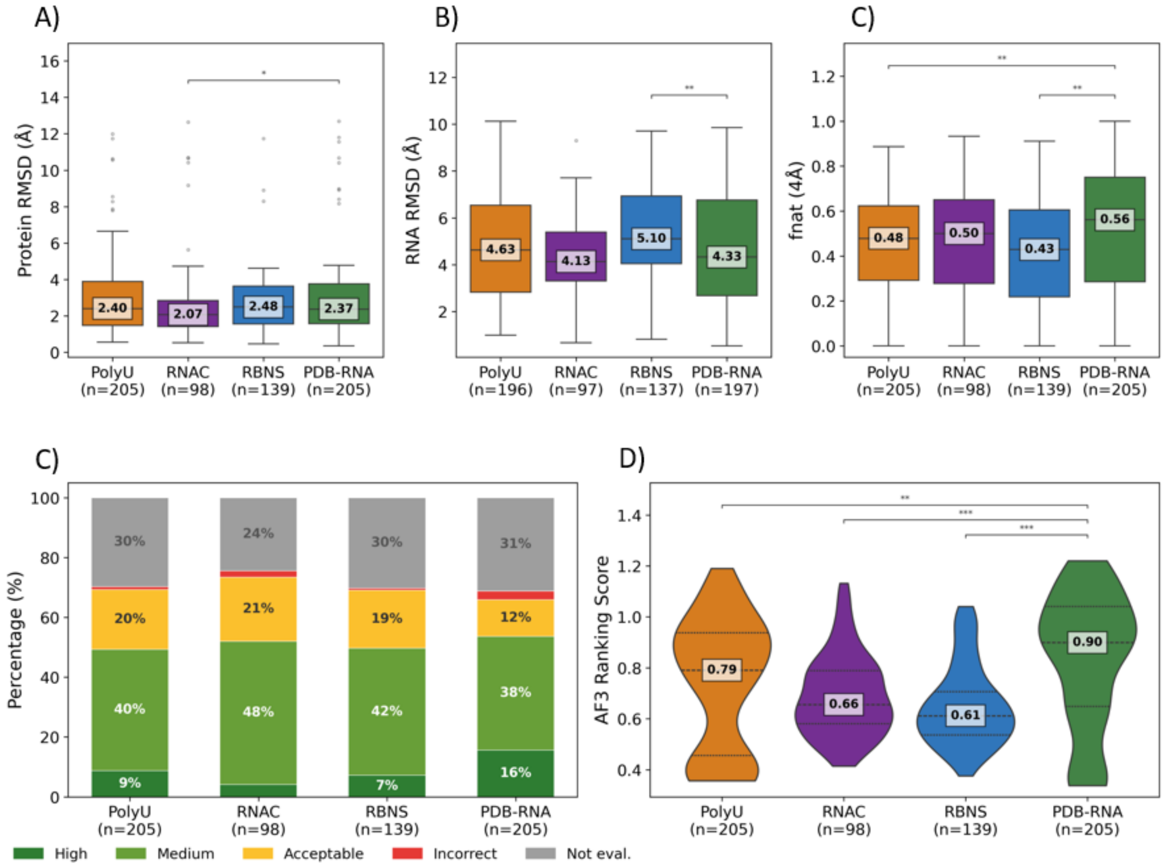


***Figure S4: AlphaFold3 specificity analysis (as in main Figure 3) when removing SNRPA-related interfaces from the dataset. A)*** *Boxplots showing the RMSD of the protein backbone for the top-ranked AF3 models obtained using four RNA input types.* ***B)*** *Same as panel A, for the RNA RMSD.* ***C)*** *Boxplots showing the fraction of native contacts (fnat) for the top-ranked AF3 models obtained using four RNA input types.* ***D)*** *Systematic evaluation of AF3 models for all conditions using CAPRI metrics.* ***E)*** *AF3 ranking scores for top-ranked models in each of the four conditions.*

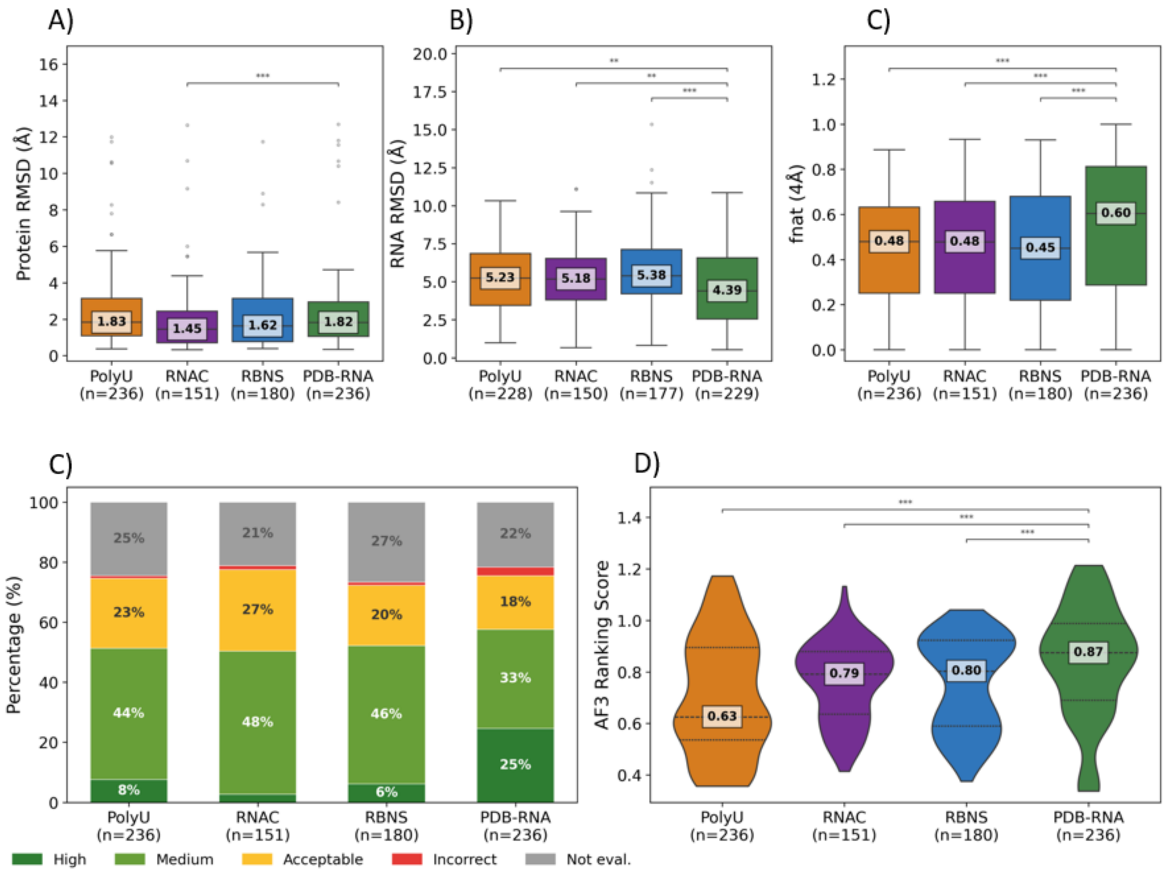


***Figure S5: AlphaFold 3 specificity analysis (as in main Figure 3) after applying technique-specific agreement score filtering (S > -2.25). A)*** *Boxplots showing the RMSD of the protein backbone for the top-ranked AF3 models obtained using four RNA input types.* ***B)*** *Same as panel A, for the RNA RMSD.* ***C)*** *Boxplots showing the fraction of native contacts (fnat) for the top-ranked AF3 models obtained using four RNA input types.* ***D)*** *Systematic evaluation of AF3 models for all conditions using CAPRI metrics. E) AF3 ranking scores for top-ranked models in each of the four conditions.*

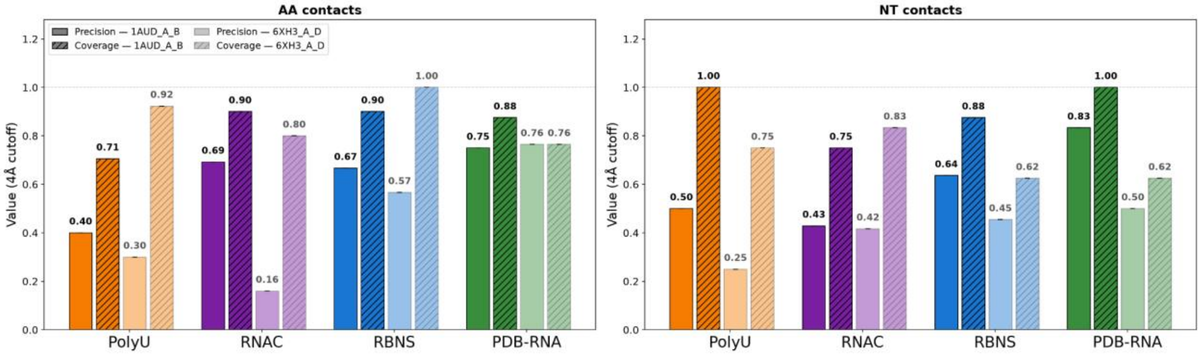


***Figure S6:*** *Precision and coverage for PolyU, RNAC, RBNS, PDB RNA conditions with 1AUD and 6XH3 interfaces.* ***A)*** *based on amino acids* ***B)*** *based on nucleotides.*

### Supplementary Tables

| Motif core | n (RNAC) | S (RNAC) | n (RBNS) | S (RBNS) |
|---|---|---|---|---|
| **GCAC** | **68** | **-0.49** | **68** | **-0.42** |
| CACA | 1 | -0.69 | — | — |
| NCAC | — | — | 1 | -0.76 |
| AGCN | 2 | -1.62 | — | — |
| GCNN | — | — | 2 | -1.73 |
| GAGC | 4 | -2.44 | — | — |
| AGCU | — | — | 4 | -2.79 |

***Table S1: Motif cores identified by the agreement scoring workflow for SNRPA**. All identified motif cores in RNAC and RBNS are listed, with the number of associated interfaces (among 75 structures involving SNRPA) and associated agreement score.*

| Structure | PDB-RNA | PolyU | RNAC | RBNS |
|---|---|---|---|---|
| **1AUD** | 0.99 | 0.60 | 0.93 | 0.89 |
| **6XH3** | 0.76 | 0.57 | 0.85 | 0.95 |

***Table S2:** AlphaFold 3 ranking score for 1AUD and 6XH3 five sampled models using different RNA sequence: from the experimental structure (PDB-RNA), from RNAcompete binding profile (RNAC), for RNA-Bind-N-Seq binding profile (RBNS) and a polyU.*

```
gagga---
-aggug--
-aggag--
-cggag--
-uggag--
--gguga-
---aaggu
--ggagu-
--gaagg-
---gaggu
---gagaa
--ggaga-
--ggagg-
```

***Table S3***: *Alignment of LIN28A RNA 5-mers obtained from RNAC binding preference data.*

| Structure | PDB-RNA | PolyU | RNAC |
|---|---|---|---|
| **5UDZ** | 0.96 | 0.53 | 0.53 |
| **2LI8** | 1.01 | 0.72 | 0.76 |

***Table S4***: *AlphaFold 3 ranking score for 5UDZ and 2LI8 five sampled models using different RNA sequence: from the experimental structure (PDB-RNA), from RNAcompete binding profile (RNAC) and a polyU.*
